# Antisense transcriptional interference mediates condition-specific gene repression in budding yeast

**DOI:** 10.1101/169730

**Authors:** Alicia Nevers, Antonia Doyen, Christophe Malabat, Bertrand Néron, Thomas Kergrohen, Alain Jacquier, Gwenael Badis

**Affiliations:** Unité GIM, Institut Pasteur, Paris, France; CNRS UMR3525, Paris, France; Université Pierre et Marie Curie, Paris, France; Bioinformatics and Biostatistics Hub, C3BI, Institut Pasteur, USR 3756 IP CNRS, Paris, France

## Abstract

Pervasive transcription generates many unstable non-coding transcripts in budding yeast. The transcription of such noncoding RNAs, in particular antisense RNAs (asRNAs), has been shown in a few examples to repress the expression of the associated mRNAs. Yet, such mechanism is not known to commonly contribute to the regulation of a given class of genes. Using a mutant context that stabilised pervasive transcripts, we observed that the least expressed mRNAs during the exponential phase were associated with high levels of asRNAs. These asRNAs also overlapped their corresponding gene promoters with a much higher frequency than average. Interrupting antisense transcription of a subset of genes corresponding to quiescence-enriched mRNAs restored their expression. The underlying mechanism acts in *cis* and involves several chromatin modifiers. Our results convey that transcription interference represses up to 30% of the 590 least expressed genes, which includes 163 genes with quiescence-enriched mRNAs. We also found that pervasive transcripts constitute a higher fraction of the transcriptome in quiescence relative to the exponential phase, consistent with gene expression itself playing an important role to suppress pervasive transcription. Accordingly, the *HIS1* asRNA, normally only present in quiescence, is expressed in exponential phase upon *HIS1* mRNA transcription interruption.

## INTRODUCTION

In steady state, the transcriptome reflects the equilibrium between RNA synthesis and degradation. Eukaryotes have developed sophisticated systems to control the turnover of mRNAs and ncRNAs necessary to the cell, undesired RNA species being rapidly eliminated by quality control mechanisms.

The development of genome-wide techniques such as tiling arrays and cDNA next-generation sequencing to analyse transcriptomes revealed that eukaryotic genomes are pervasively transcribed (1). The genome of budding yeast is particularly compact and it has been hitherto conceded that more than 70% of it is composed of protein coding ORFs (2). Yet, this is only true if one does not distinguish the two DNA strands. If one takes into account sense and antisense genomic DNA, non protein-coding sequences represent up to 65% of it, leaving room to a large fraction of the genome for the generation of pervasive non-coding transcripts.

In yeast, pervasive transcription has been first reported more than a decade ago. If a fraction of it was uncovered in wild-type cells (3, 4), a substantial part of the eukaryotic pervasive transcription is “hidden” as it generates very short-lived “cryptic” transcripts. These RNAs are difficult to detect unless they are stabilized by interfering with quality control mechanisms that normally eliminate them (5). Pervasive transcripts detected in wild-type yeast cells have been named “SUTs” for “Stable Unannotated Transcripts” (4), and different names have been given to cryptic transcripts depending on which factor was mutated in order to stabilise a particular class of RNAs. For example, CUTs (Cryptic Unstable Transcripts) were characterised upon removal of Rrp6, an exonuclease specific of the nuclear form of the exosome (4, 6, 7), XUTs were revealed upon removal of the cytoplasmic exonuclease Xrn1 (8) and NUTs correspond to transcripts that accumulate when the nuclear termination factor Nrd1 is depleted (9). Yet, there are in yeast only two main pathways responsible for the efficient elimination of pervasive transcripts: the nuclear Nrd1-Nab3-Sen1 (NNS) pathway, in which the early transcription termination of cryptic transcripts by the NNS complex is coupled to the degradation by the nuclear TRAMP-exosome complex (9–12) and the cytoplasmic non-sense mediated mRNA decay (NMD) pathway (13, 14). Many of pervasive transcripts require both pathways for their efficient and fast elimination (see 13).

Irrespective of which pathway predominates for their degradation, these transcripts all originate from nucleosome free regions (NFRs), which are essentially found 5’ and 3’ of mRNA coding sequences (15). When they originate from 5’ NFRs, they are most often transcribed divergently from mRNAs and result from an intrinsic low polarity of gene promoters (4, 7). This divergent transcription has the potential to interfere with the expression of the neighbouring upstream gene. Likewise, when a noncoding transcript initiates from the 3’ NFR in an antisense orientation to the upstream gene, its transcription has the potential to interfere with the proper expression of the corresponding mRNA (8, 16). Such transcription interference by pervasive transcription is largely prevented genome-wide by the NNS quality control pathway, which ensures the early transcription termination of these transcripts and prevent them to extend into the promoter region of the corresponding antisense genes (9–11, 17).

Whether pervasive transcription has a general function is a matter of debate. The fact that highly efficient quality control mechanisms have been selected during evolution to eliminate most of these transcripts argue in favour of the idea that most of them are non functional; however pervasive transcription by itself, more than its product, could play a role. Yet, the existence of the NNS pathway, which, by terminating pervasive transcription early, is key in preserving pervasive transcription from interfering with the expression of many coding genes, also suggests that a large fraction of these events simply result from the low specificity of RNA polymerase II (PolII) transcription initiation.

There are a number of well-documented examples of individual coding gene regulation through the transcription of a non-coding RNA: *SER3* (18), *IME1* and *IME4* (19), *GAL10/GAL1* (20, 21), *PHO84* (22), *CDC28* (23) as examples. In the vast majority of cases analysed in budding yeast, the synthesis of a non-coding transcript has only an effect in *cis*. The prevailing model is that repressive chromatin marks are deposited in the promoter regions of genes in the wake of RNA polymerase II (PolII) transcribing the associated non-coding RNAs (24, 25). It is thus the act of transcription rather that its product, which is important. Several distinct mechanisms can be at play, but the general theme is that methyltransferases interacting with the the carboxy-terminal domain (CTD) of the PolII large subunit deposit histone methylation marks that recruit repressive chromatin modifiers such as histone deacetylases or nucleosome remodelling complexes. In budding yeast, there are two such CTD associated histone methyl transferases. Set1 methylates histone H3K4 at promoters and gene proximal regions of actively transcribed genes while Set2 methylates H3K36 at more distal gene regions. The role of Set1 is complex. It is responsible for both H3K4 di- and tri-methylation (H3K4me2 and H3K4me3). It has been proposed that H3K4me3 at the beginning of actively transcribed genes could enhance and help maintaining pre-initiation complex assembly and an active acetylated chromatin state, thus playing a positive role on transcription. Conversely, Set1 generates H3K4me2 in the body of gene, which recruits the histone deacetylase complexes SET3 or RPD3L, resulting in transcription initiation repression (see 26 for review). Set2 is responsible for the H3K36 methylation (H3K36me2) in the body of genes, resulting in the recruitment of the Rpd3S deacetylase complex that plays an essential role in preventing improper internal initiation (27, 28). Thus both Set1 and Set2 have the potential to mediate transcriptional interference and have been implicated in gene repression by non-coding RNA transcriptional interference (see 24 for review).

Does pervasive transcription, and in particular antisense transcription, play a larger role in gene regulation? If so, the act of transcription by itself may constitute a critical step in that pathway. If not, apart from a few exceptions, pervasive transcription may only represent transcriptional noise.

Several large-scale studies attempted to answer this question. Genes with large expression variability (such as stress response and environment specific genes) often have antisense expression suggesting a general regulatory effect of antisense on gene expression (29). Others correlated antisense expression with chromatin marks, either in a wild-type context or with a *rrp6* mutant (17, 25, 30, 31) but no global anti-correlated trend was found between asRNA and mRNA expression.

Very recently, to which extent asRNA transcription could act on gene regulation was examined by measuring, under various conditions, the effect of specific antisense SUTs transcription interruption on the expression of the corresponding proteins fused to GFP (32). This study showed that, for 12-25% of genes associated with an antisense SUT, a detectable but weak antisense-dependent gene regulation could be observed under at least one condition. Although no specific biological pathway seemed enriched in the tested asRNA responsive genes, the analysis showed that repression by asRNA transcription interference helps reducing somehow mRNA expression basal levels, especially for genes expressed at a low level, reinforcing complete gene shut off. However, the analysis was restricted to SUTs, *i.e.* non-coding RNAs readily detectable in wild-type cells, which are limited compared to the reality of antisense transcription in the cell as we know that SUTs represent only a minority of the pervasive transcripts, most of which are too unstable to be detected in wild-type cells (4, 7, 8, 33).

The nuclear NNS quality control pathway prematurely terminates the transcription of many of the pervasive RNAs to prevent them from interfering with mRNA expression (9, 34). However, many pervasive transcripts escape, at least in part, this first surveillance pathway and are extended up to cryptic cleavage and polyadenylation sites (polyA sites), potentially over the transcription start site (TSS) of their associated genes. This can lead to the export of long non-coding RNAs into the cytoplasm, where they are rapidly degraded by the NMD pathway (13, 14).

In order to measure a relevant “antisense transcriptome”, we analysed genome-wide the amount of asRNAs associated to each mRNA in a NMD mutant context (*upf1*Δ). In this mutant, hidden pervasive transcripts that escaped the nuclear NNS surveillance accumulate in the cytoplasm and can thus be quantified. An important fraction of the less expressed genes are associated with asRNAs, especially if the asRNAs overlapped the associated sense gene promoter. In addition, many of these genes with promoter-overlapping asRNAs were enriched for genes up-regulated in chromatin remodelling mutants such as *set2*Δ or *set1*Δ. Interestingly, the majority of mRNAs enriched during the stationary phase (G0) fall in the category of genes poorly expressed during the exponential phase and 30% of them are associated with antisense RNAs overlapping their promoter, a much higher proportion than overall average (9.5%). These observations strongly suggest that this particular class of genes is frequently subjected to asRNA transcription interference for full repression during exponential growth, a prediction we validated experimentally for a subset of genes.

Our study showed that antisense-mediated transcriptional interference is, in budding yeast, a mechanism more frequently used than anticipated when mRNA expression needs to be tightly repressed under specific conditions.

## MATERIAL AND METHODS

### Yeast strains and cultures

All strains are listed in Supplementary Table S1, are derivative of BY4741 or BY4742, and were obtained from the Euroscarf deletion collection (http://www.euroscarf.de/). A 37 nucleotides sequence constituting the NNS terminators (GTAATGAATTAAGTCTTGATATATAACAATTAGCTTG construct 78-wt in (35)) was inserted into BY4741 or BY4742 strains using the seamless cloning-free PCR-based allele replacement methods as described in (36).

Briefly, gene-specific PCR products containing adaptamer A or adaptamer B and NNS terminator were reconstituted with two successive PCR using A-GENE primer and GENE_NNS_S/AS_rev (PCR1) and GENE_B and NNS_AS_GENE_fwd (PCR2), followed by A_GENE and GENE_B (PCR3). GENE stands for ARO10, PET10 NNS_S, SHH3 MOH1 and CLD1 (see Supplementary Table S2). In parallel Fragment L and R were obtained using primer CS1199/CS1200 and CS1201/CS1202 on a URA3 K. lactis DNA template from plasmid pBS1539, (37). All PCR were done with a high-fidelity Phusion^®^ HighFidelity (NEBiolabs), following the manufacturer’s instructions.

For PET10_NNS antisense, a PCR product GB988/GB989 obtained using pFL38 (from http://seq.yeastgenome.org/vectordb) as a DNA template was used to transform BY4741 plated on SC-URA medium. [URA3+] clones were transformed with 100 pmol of annealed GB990/GB991 primers, plated on YPD at 30°C overnight, and replicated on 5FOA medium in order to select *URA3* popped-out constructs. All the constructs were sequence-verified.

Supplementary Figure S1 lists the position of NNS terminator insertion, in both sense and antisense orientation. Strains and oligonucleotides are listed in Supplementary Table S1 and Table S3 respectively.

Cells were grown to mid-exponential phase in YPD-rich medium at 30 °C, and homogeneous populations were purified as “Quiescent cells” (or G0), obtained from a stationary phase culture after 10 days of growth at 30°C in YPD and purification of the dense fraction on percoll gradient according (38).

#### RNA extraction

Total RNA from logarithmic and G0 cells were extracted with guanidium thiocyanate phenol-chloroform following (39) with the addition of 500 μl of glass beads prior to solution D addition, and vortex in a MagNA lyser (Roche) 90 seconds at 4800 rpm after solution D addition.

#### Libraries preparation

3’ Long SAGE libraries were constructed as described in (40), except than total RNA were extracted from BY4741 logarithmic and G0 cells using the guanidium thiocyanate phenol-chloroform procedure described in (39).

TruSeq stranded mRNA LT sample prep kits (Illumina) were used to prepare RNAseq libraries, on RiboZero gold (Illumina) treated RNA according the manufacturer’s instruction. Single read 50 (SR50) sequencing were performed on an Illumina Hiseq 2500 (Pasteur Transcriptomic Platform PF2).

#### Northern blot

Northern blots were carried out on 4 μg Total RNA as described in (7) using strand specific 32P-labeled riboprobes (see Supplementary Table S3) except for *SCR1* for which a 32P-labeled oligonucleotide was used (GB987).

#### Strand-specific RT-qPCR

Turbo DNase-treated RNA (Ambion) from exponentially growing yeast cells was used after acid Phenol Chloroform purification as an input for reverse transcription using 2 pmol of each gene-specific primers and 1 μg RNA using 0,5 μl of SuperScript III reverse transcriptase (Invitrogen) according to the manufacturer’s instructions but supplemented with 20 mg/ml actinomycin D (Thermo Fisher) to ensure strand specificity of the reverse transcription. For *PET10* and *SHH3* strand specific reverse transcription, a mix of 2 μM FF3033, AC407, AC429, AC500 and GB1038 primers was used for sense-specific, and FF3033, AC430, AC431 and AC63 primers for antisense-specific measurement (see Supplementary Table S3 for gene correspondence). For qPCR, cDNA samples and -RT controls were diluted 10 times, and 2μl were amplified using the qPCR Mix 2X Lo-Rox (Eurobiogreen). *CPS1* mRNA was used as the reference gene as its level does not change between exponential and G0 phases.

### Data analysis

#### Illumina reads treatments

For RNAseq libraries, duplicated reads were first filtered out using fqduplicate (ftp://ftp.pasteur.fr/pub/gensoft/projects/fqtools/fqtools-1.1.tar.gz). Then sequencing error were corrected using Musket (41; version 1.1). Reads of bad quality were removed using fastq_qual_trimmer (https://github.com/ivars-silamikelis/fastq_qual_trimmer, version 1.0) with a threshold of 20. Illumina adpaters were finally removed using Flexbar (42; version 2.7). After removal of the random sequence tag, resulting reads were mapped using bowtie (43; version 2.2.3 with the following parameters: –N 1 –p 1 —no-unal –D 15 –R 2 –L 22 –I S,1,1. 15) and a compilation of *S. cerevisiae* genome (S288C reference sequence, Release 64 obtained from the Saccharomyces Genome Database (SGD) [http://www.yeastgenome.org/]) and *S. pombe* genome (ASM294 reference sequence, v2.19 obtained from PomBase [http://www.pombase.org/]) as reference genomes.

For 3’Long SAGE libraries, duplicated reads were first filtered out using fqduplicate. Illumina adpaters were then removed using AlienTrimmer (44). Reads corresponding the 3’ end of transcripts were identified by detection of a polyA sequence at the end of the reads with a minimal size of 6 nucleotides. After Poly A removal, the resulting reads were mapped using bowtie (same version and parameters that above) and the *S. cerevisiae* genome (S288C reference sequence, Release 64 obtained from the Saccharomyces Genome Database (SGD) [http://www.yeastgenome.org/]). False positive reads (*i.e.* reads identified by a ≥ 6nt encoded polyA sequence but not a true 3’end) were filtered out by matching with encoded PolyA sequence in the genome.

#### Mapped reads processing

For 3’ Long SAGE libraries the 3’ -end positions of the resulting mapped reads were used as TTS positions and extracted to wig files. For RNAseq libraries, reads corresponding to the whole transcripts and full read coverage were extracted to wig files.

#### Normalisation and differential expression

Transcript differential expressions were calculated using DESeq2 (4) within the SARTools pipeline (45; version 1.4.1).

Sample corresponding to cells in exponential phase were first treated together as a separated group, as well as samples corresponding to cells in G0. During the process, SARTools performed a normalisation step. Normalisation factors were extracted and used to produce normalised wig files.

G0 samples were normalised in a second time against exponential phase sample using the spike-in of *S. pombe* transcripts. *S. pombe* transcripts median reads counts were determined for each sample after the first normalisation step. Then a global mean for *S. pombe* transcripts reads counts was calculated for quiescent and exponential phase samples. A ratio Exponential/Quiescent was calculated and applied to all G0 samples (wig files and transcripts reads counts).

#### Heatmap counting and visualisation

Antisens / mRNA coverage was counted and visualized in a −50-+200 nucleotides windows using the Counter RNAseq window (CRAW) package version 0.9.0 (https://pypi.python.org/pypi/craw/0.9.0).

## RESULTS

### Characterisation of antisense transcription in *upflΔ* cells

In order to reveal antisense transcription that escaped the nuclear NNS surveillance pathway, we quantified the asRNAs levels in the proximal region of the protein coding genes using a +1 (mRNA TSS) to +200 nucleotides window with a strain impaired for the cytoplasmic NMD surveillance pathway (*upf1*Δ mutant). Figure 1A shows these values (y axis) plotted against the average mRNA levels per gene (number of reads per nucleotide; x axis). Similar to previously reported data (25), a linear regression analysis did not reveal any correlation between the levels of antisense transcription and that of the corresponding mRNAs (Pearson correlation coefficient R^2^=0.07). Yet, the less expressed mRNAs appeared to be generally associated with high levels of asRNAs. In order to quantify this observation, we partitioned the genes according to their mRNAs levels in ten bins with an equal numbers of genes. The less expressed genes (bin 1) had significantly higher levels of asRNAs than the genes within higher mRNA expression categories (bins 2 to 10; see Figure 1B, Dataset 1 and Supplementary Table S3).

**Figure 1.**
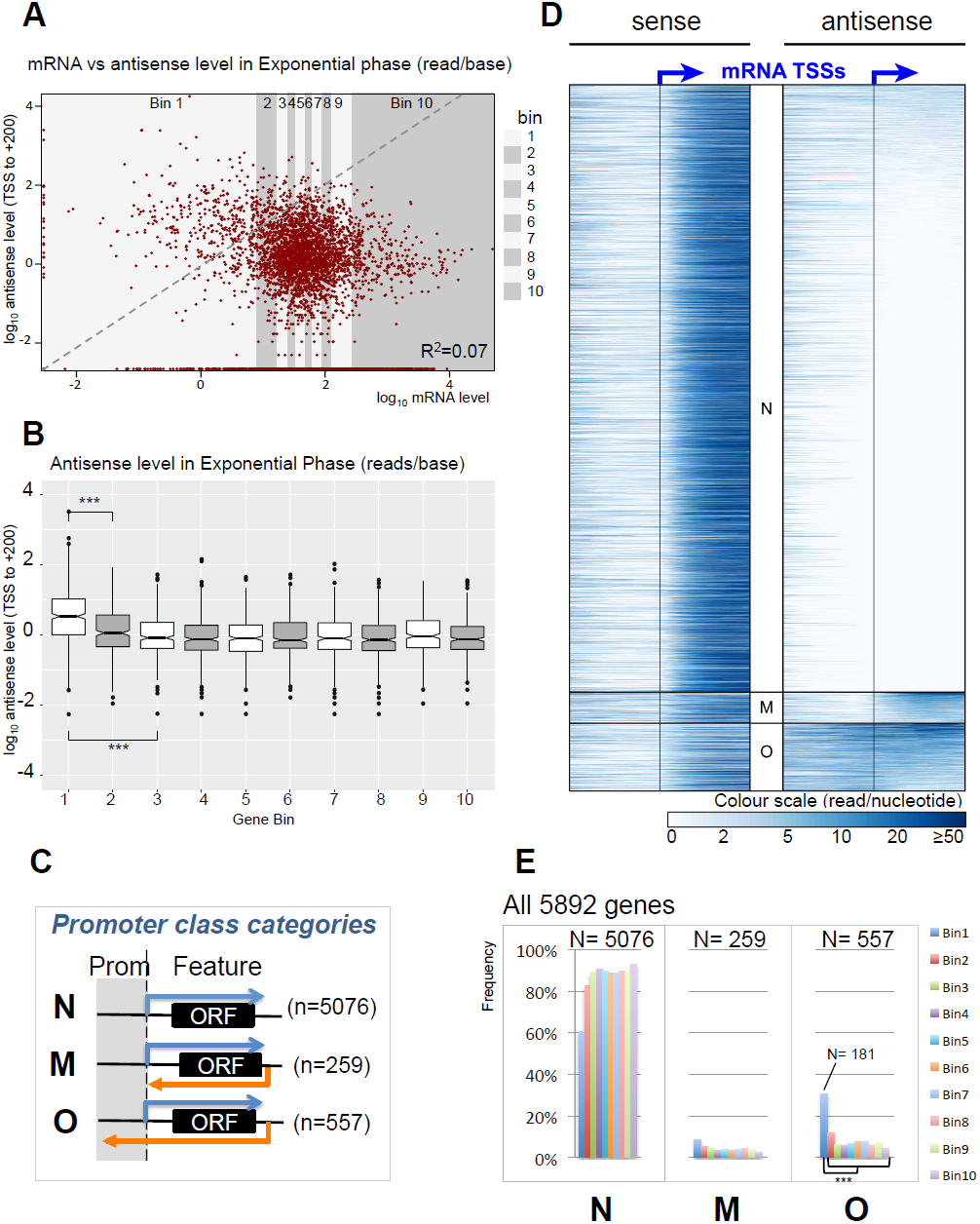
Antisense ncRNAs are over-represented in lowly expressed genes. A Scatter plot representing the antisense level (ordinate) function of the corresponding mRNA level (abscissa) in log_10_ read/base. Gene read count was determined for 5892 genes, and divided into ten bins (grey strips) of equal length (N = 589 genes per bin for bin1 to bin9; N=591 genes for bin10). The Pearson correlation coefficient R^2^ = 0.07. B Comparison of the average antisense level distribution between bins. Boxplots shows the distribution of the average antisense levels within each bin. Brackets indicate the results of an Anova test on pairs of distributions, with *** for P < 0.001. C Schematic of the gene-associated promoter class categories depending of the presence and the characteristics of asRNA: N = No asRNA, M = asRNA within the mRNA, O = TSS-overlapping asRNA. An arbitrary threshold of at least three RNA sequencing reads per nucleotide, in a +1 to +200 nucleotide window relative to the mRNA TSS position, was used to define the presence of an asRNA. D Heatmap distribution of mRNA and antisense around the TSS of all genes, sorted by antisense and promoter class categories. Depending the class of promoter defined in C, a category N, M or O was assigned to each gene. E Promoter class categories count per bin. The total number of genes that belong to each class of promoter is indicated (Class N: N=5076; Class M: N=259; Class O: N= 557). Bar charts represent the percentage of each class within the 10 bins defined in A (see also Dataset 1). Brackets indicate the results of a statistical inference test on pairs of distributions between bin1 and each other bins, with *** for P < 0.001.

At least two non-exclusive phenomena could explain this observation. First, gene transcription itself could have a repressive effect on asRNA transcription initiation from their corresponding gene-3’ NFRs (29). Hence, asRNAs initiating within NFRs situated downstream of non-expressed genes should be less subjected to such repression by coding gene transcription. Conversely, antisense-transcription from 3’ NFRs could be a common mean to contribute to a tight gene repression. If the former explanation is correct, asRNAs associated with non-expressed genes should not show different termination characteristics than other asRNAs. In contrast, it was shown that repression by asRNAs correlates with mRNA TSS overlap (32). If asRNAs associated with the less expressed genes contribute to their tight repression, these asRNAs should overlap the mRNA TSSs more often than other asRNAs. We thus categorized genes depending on the occurrence of their associated asRNAs across TSSs by analysing a window between -50 nucleotides to +200 nucleotides relative to the mRNA TSS. We defined three types of genes. Genes without substantial asRNAs over the +1 to +200 nucleotide region (arbitrarily set below three reads per base over this window) defined class N (No antisense). Genes with asRNAs but with an average read number below three in the −50 to −1 nucleotide region, thus terminating before the mRNA TSS, defined class M (mRNA antisense). Conversely, genes with asRNAs with an average read number above three in the -50 to -1 nucleotide region defined genes with TSS overlapping asRNAs (class O - Overlapping antisense) (Figure 1C; Dataset 1). Figure 1D shows a heat map of the sense and antisense transcripts over a −200 to +200 nucleotides window around the mRNA TSSs, classified according to the three classes. Among the 5892 protein coding genes analysed, 5076 belong to class N, 259 to class M and 557 have an overlapping asRNA (class O). The higher proportion of asRNAs in bin 1 mostly resulted from the over-representation of class O asRNAs, which represent 30% of bin1 (181 class O genes among the 590 genes in bin1). Class O asRNAs represented 78% of all asRNAs of bin 1 (181/233), while this proportion was only of 64.5% (376/583) within all the other bins (Figure 1E, Supplementary Figure S2 and Dataset 1). It strongly suggested that the higher number of asRNAs associated genes in bin 1 relative to the others bins reflected a potential regulatory role associated with a number of these asRNAs.

### Genes up-regulated in the absence of chromatin regulators are enriched in the class of poorly expressed genes with TSS-overlapping asRNAs

If antisense transcription can affect sense transcription, one should expect that genes associated with asRNAs be more up-regulated in chromatin modifier mutants implicated in transcriptional interference, in particular *set1* and *set2* mutants. Given that antisense transcriptional interference involves the extension of asRNA up to the promoter regions of the genes, Set2, which promotes H3K36me2 at late stages of PolII elongation, seemed a good candidate to mediate gene repression by asRNA transcription. *SET2* mutants are intrinsically difficult to analyse by RNAseq or tilling arrays since a major role of Set2 is to suppress both sense and antisense internal initiation within gene transcribed regions (13, 27, 28, 46). The cryptic initiation events observed in *set2*Δ mutants in the sense orientation can thus lead to misleading quantitation due to the overall increase of sense RNAseq counts (13). We thus took advantage of the analysis of individual TSSs in the Malabat *et al.* study, which allows the quantitative analysis of the specific mRNA TSSs, irrespective of internal transcriptional initiation. We considered a gene as up-regulated upon *SET2* deletion when its strongest mRNA-linked TSS cluster was induced at least two fold with a p-value ≤0.05 (Supplementary file 3 in 13). Ninety-five of 5228 genes analysed in this dataset were up-regulated in a *set2*Δ *strain* (see Dataset 1) Strikingly, genes with TSS-overlapping asRNAs (class O) showed the highest percentage of up regulation in a strain lacking *SET2* (9,1% compared to 1.8% for all genes; Figure 2A). Combining gene promoter classes with the mRNA expression level categories drastically increased this bias since class O of bin 1 showed the highest proportion of genes up-regulated in *set2*Δ cells (Figure 2B, right panel).

**Figure 2.**
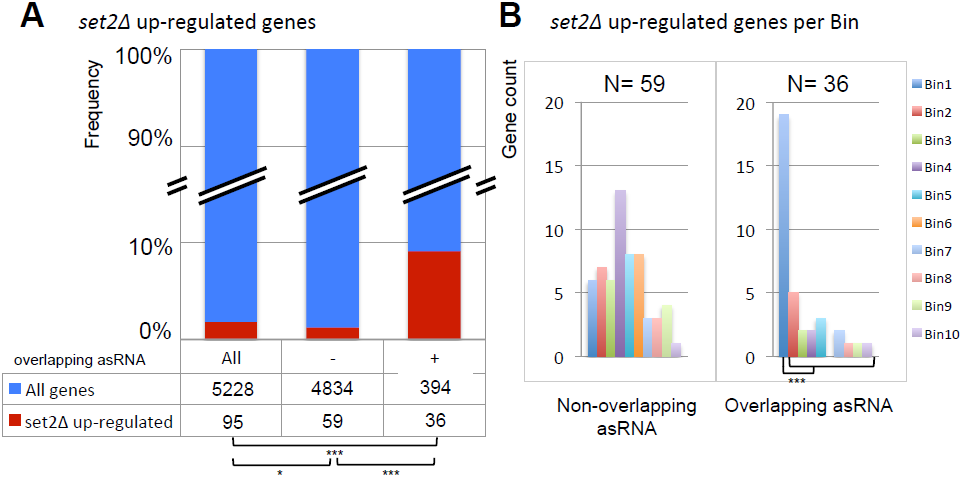
Promoter overlapping antisenses are overrepresented in *set2*Δ targets. A Gene distribution across promoter categories in *set2*Δ up-regulated genes (dataset from 13). Stacked histograms represent the proportion of *set2*Δ up-regulated genes across all genes (All), or depending the presence of a TSS-overlapping asRNA (“+” = class O) or not (”−” = classes N+M). Brackets indicate the results of a statistical inference test on pairs of distributions, with * for P < 0.05, and *** for P < 0.001. B *set2*Δ up-regulated genes count depending the presence of a TSS-overlapping asRNA or not and per bin. Brackets indicate the results of a statistical inference test on pairs of distributions between bin1 and each other bins, with *** for P < 0.001

Direct measurement of transcription levels by NETseq have been analysed in a *set2*Δ mutant (47). Although a higher number of genes were found to be up-regulated in absence of Set2, possibly due to internal initiation events not being filtered out, the same trend was observed (Supplementary Figure S3A). This prompted us to analyse the data for the *set1*Δ, as well as *rco1*Δ and *eaf3*Δ (two components of the Rpd3S deacetylase complex) mutants from the same dataset, as these factors have also been found to be involved in transcriptional interference. Genes up-regulated upon deletion of these genes were also clearly over represented in class O (Supplementary Figures S3B-D). Altogether these results suggest that repression by antisense transcriptional interference is frequent for poorly expressed genes, a process mediated by several chromatin-modifying factors linked to elongating PolII. Supplementary Figure S3E reports the large number of up-regulated genes overlapping in the different mutant strains, which highlights the redundancy of these processes (47).

### Quiescence enriched genes are associated with high levels of asRNAs

We next determined if poorly expressed genes (bin 1) belong to a particular category of regulated genes. An expected category of genes strongly repressed during exponential growth are those found enriched in stationary phase and/or in quiescence (G0). We thus analysed a dataset reporting the time course of mRNA expression of a wild-type strain over a complete 10-days growth. Figure 3A shows that the stationary phase-enriched genes (SP-enriched in 48) are the most abundant in bin 1. However in stationary phase, the cell population might not be homogeneous since it is composed of dead, senescent and quiescent cells (38, 49). To circumvent this problem, we performed a genome-wide RNAseq analysis using a homogenous population of quiescent cells derived from wild-type or *upf1*Δ strains in order to analyse both gene and pervasive transcription (see Material and Methods). To normalize the overall level of transcripts per genome, we spiked in the budding yeast cultures before RNA extraction with identical reference aliquots of a *Schizosaccharomyces pombe* exponential culture (see Materials and Methods for the normalization procedure). We defined quiescence-enriched (Q-enriched) mRNAs as being, following normalization, five times more abundant in the G0 population than the exponential growing phase (total of 261 genes, Supplementary Table S4). As anticipated, Q-enriched mRNAs were found in majority within bin 1 (163 genes in bin1 among the 261 Q-enriched genes; Figure 3B and C). Accordingly, Figure 3D shows that, as for genes within bin 1, Q-enriched mRNAs were associated with higher asRNA levels than random (*** with p = 1.5 10^-4^) and their distribution in the different asRNA associated genes classes (classes N, M & O) was similar to that of bin 1 (Supplementary Figure S4A). While, upon deletion of *SET2,* 1.8% of all genes are up regulated, this fraction rises to 9.4% of all genes with TSS-overlapping asRNAs (Figure 2A) and to more than 35% when only considering Q-enriched genes (Supplementary Figure S4B). Breaking down these figures by bins and promoter classes showed that this strikingly high proportion was primarily contributed by class O genes, representing 13 out of 22 (59%) of the *set2*Δ up-regulated Q-enriched genes (Figure 3E). These observations strongly suggested that asRNA transcriptional interference could be a frequent mechanism of tight repression for this specific class of genes. In order to directly test this hypothesis, we chose for further analysis five representative examples of Q-enriched genes associated with an asRNA spanning the mRNA TSS: *PET10*, *SHH3*, *MOH1*, *CLD1* and *ARO10*. Among these genes, only *ARO10* was previously tested for asRNA mediated transcription interference (32).

**Figure 3.**
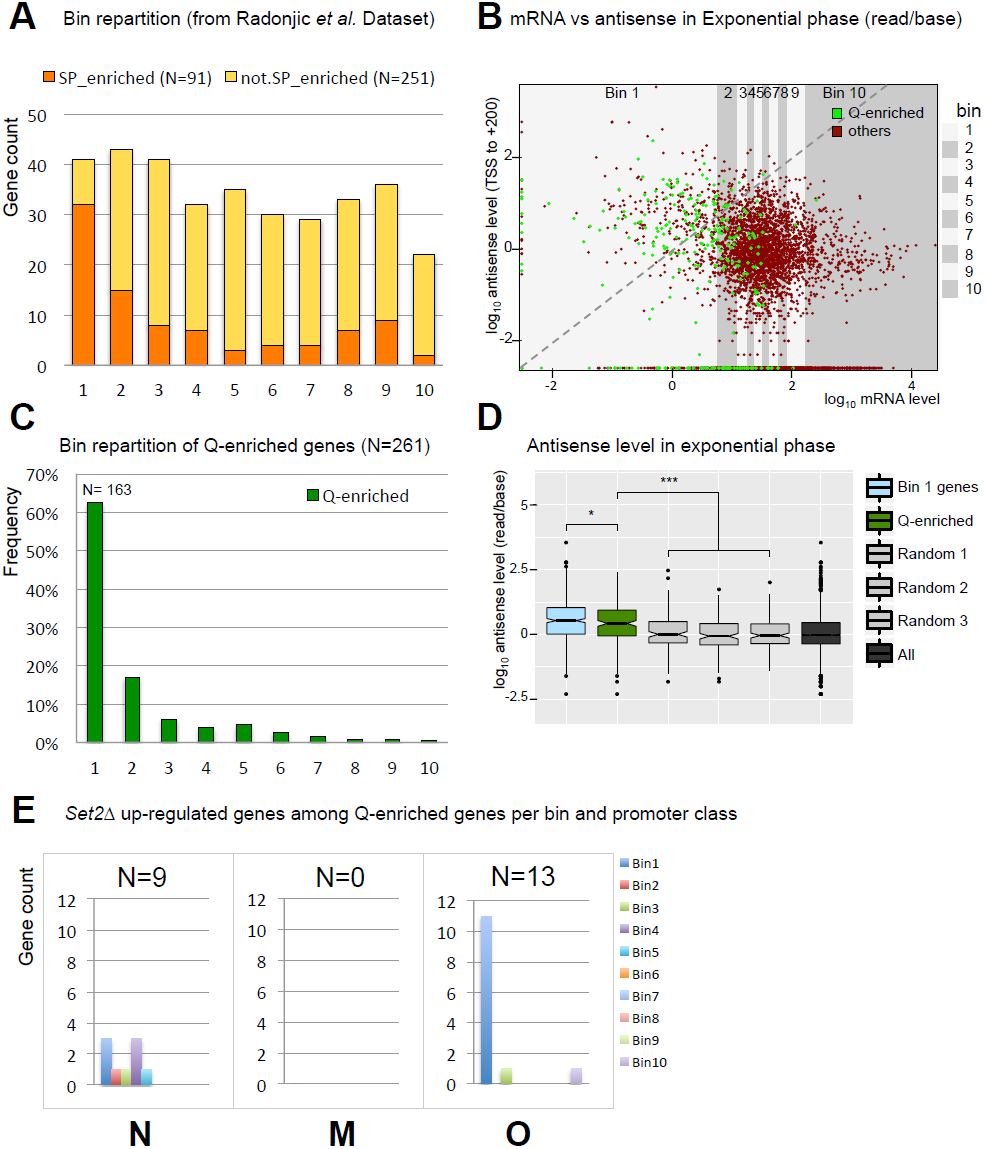
Quiescent-enriched genes are associated with high antisense level. A Bar plot of stationary phase-enriched genes count (SP-enriched) versus other genes count (not SP-enriched) within each bin (dataset from 48). B Distribution of quiescence-enriched genes among the 5892 yeast genes. Scatter plot of the antisense level as a function of the corresponding mRNA level. 261 genes were found enriched at least 5 times between exponential and quiescence, defining the quiescence-enriched genes (Q-enriched, green dots, see also Dataset 1 and Materials and Methods). C Bar chart of the 261 Q-enriched genes within the 10 bins. D Distributions of antisense level for different gene categories in exponential phase. Boxplots show the mean antisense level of 261 corresponding Q-enriched genes (green) or “Random” (grey) genes. Random-1, -2, and -3 were defined by random sampling of 261 genes among all the 5892 genes. “Bin_1” (blue) or “All” categories (black) are the measures of all 589 genes from bin1 or all 5892 genes respectively. Brackets indicate the results of an Anova test on pairs, with * for P < 0.05, and *** for P < 0.001. E *set2*Δ up-regulated genes count among Q-enriched genes per promoter class and bin. The bar charts represent the count of *set2*Δ up-regulated genes within each category of promoter and each bin (see also Dataset 1).

### Time course of quiescence-enriched mRNAs and corresponding asRNAs show an inverse expression pattern

In order to examine the relative behaviour of these mRNAs in relation to their associated asRNAs, we performed Northern-blots time course experiments starting (t0’) by the addition of rich medium to quiescence purified cells and using strand-specific RNA probes. The five selected Q-enriched mRNAs were not only accumulating during quiescence but were in fact strongly induced after about 48 hours of culture (Figure 4), which coincides with the post diauxic shift transition (48). The asRNAs started to accumulate between 5 and 30 minutes upon rich medium addition to reach a peak of expression at ~24 hours, after which they rapidly disappeared. The mRNAs followed the inverse trend with the exception of *ARO10* that was not as substantially repressed during the exponential phase. These observations are compatible with the asRNA transcription contributing to mRNA repression. Conversely, they are also compatible with induction of the mRNA repressing the associated asRNAs.

**Figure 4.**
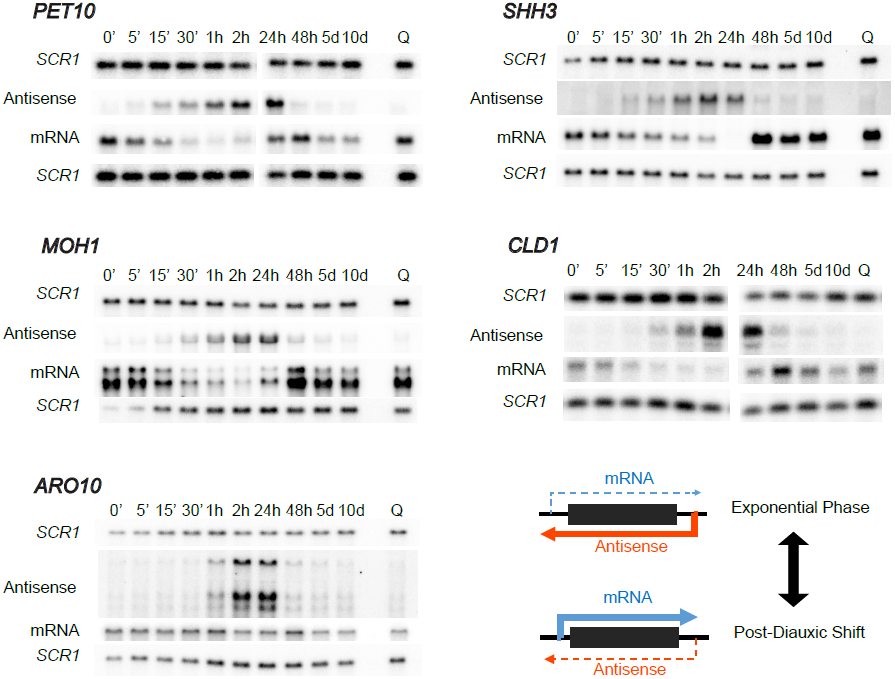
Quiescence-enriched genes mRNA and corresponding asRNAs are anti-regulated. Northern-Blot probing for time course mRNA and antisense transcripts in a Δ*upf1* strain for five examples of Q-enriched genes: *PET10*, *SHH3*, *MOH1*, *CLD1* and *ARO10*. Time point 0’ is the time at which quiescent-arrested cells are restarted in rich YPD medium. *SCR1* is used as a loading control. RNA probes are described in Supplementary Figure S1 and Supplementary Table S2.

### Interruption of antisense transcription results in de-repression of quiescence-enriched genes

One of the main effects of the NNS pathway is to prevent the expression of most pervasive transcription from interfering with the normal expression of genes genome-wide (9). This mechanism is thus intrinsically optimized to result in an early termination and in a strand specific manner. We choose to use it in order to specifically terminate asRNA transcription close to their transcription start by introducing in the TSS proximal region of the asRNAs a short (37 nucleotides) optimal NNS termination signal (NNS-ter; 35). In order to perturb as little as possible the corresponding mRNAs, we introduced this NNS-ter sequence seamlessly using a cloning-free method allowing chromosomal modifications without leaving selection markers (36). This sequence was introduced in the proximal region of the asRNAs, corresponding to the terminal region of the mRNAs (see Supplementary Figure S1 and Supplementary Table S2). The introduction of the NNS-ter signal resulted in the proper elimination of all asRNAs and in a strong up-regulation of the corresponding mRNAs, except for *ARO10* (Figure 5A). We note that *ARO10* is also, out of the five genes examined, the one that showed the weakest mRNA repression during the exponential phase (see Figure 4 and Discussion). We then verified that for NNS constructs inserted upstream the stop codon, the observed effect was not due to a NMD effect -a consequence of the ORF disruption that could insert a premature stop codon that could be recognized like a NMD substrate-but to the effect of the antisense interruption (Figure 5B lanes “if”). We also verified that the use of a scrambled, inactive version of the NNS terminator, which could not interrupt antisense transcription anymore, had no effect on mRNA expression (Figure 5B lanes “sc”).

**Figure 5.**
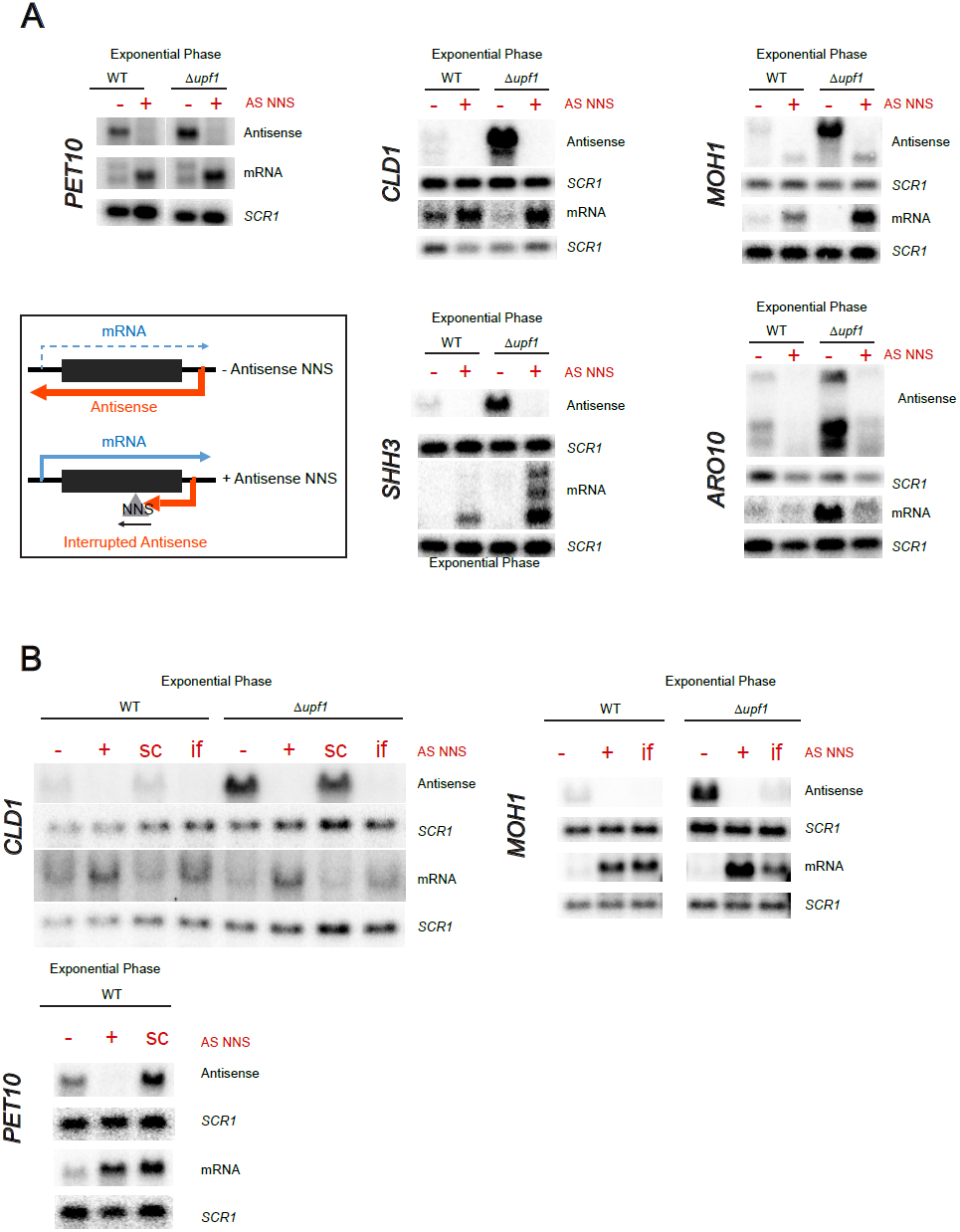
Antisense transcription interruption during the exponential growth relieves repression of quiescence-enriched genes. A Northern blot probing for the *PET10*, *CLD1*, *MOH1*, *SHH3* and *ARO10* mRNA and antisense transcripts in the WT and Δ*upf1* strains with (+) or without (−) the insertion of an antisense Nrd1-Nab3-Sen1 terminator (AS NNS). *SCR1* is used as a loading control. RNA probes and NNS insertion are described in Supplementary Figure S1 and Supplementary Table S2 (see also Material and Methods for strain construction and AS NNS-corresponding strains in Supplementary Table S1). *B* Northern blot probing for the *PET10*, *CLD1*, and *MOH1* mRNA and antisense transcripts with scrambled (sc) NNS controls (*PET10* and *CLD1*) and/or in frame (if) NNS insertion (*CLD1* and *MOH1*) (see also Material and Methods).

### The asRNA associated gene repression acts in *cis*

Although the majority of non-coding RNA associated gene regulation has been shown to act only in *cis*, a *trans* effect of the asRNA itself has been invoked in a few cases (see 24 for a review). We directly addressed this question on *PET10* by comparing the mRNA and the asRNA expression in *cis* and in *trans*. To this end, we built diploid strains where *PET10* sense and antisense transcripts were disrupted on one or two of the homologous chromosomes, allowing the expression of the asRNA either from the same chromosome as the mRNA (in *cis*), from the opposite chromosome (in *trans*) or without asRNA expression (Figure 6A). RT-qPCR measurement showed that *PET10* mRNA is repressed only when its asRNA is expressed in *cis* (blue) but not in *trans* (green). In this case the mRNA level reached the same level as observed in the control strain without antisense (red). This is fully consistent with the hypothesis that the antisense transcription and not the asRNA itself, acts to repress mRNA by a transcriptional interference mechanism.

**Figure 6.**
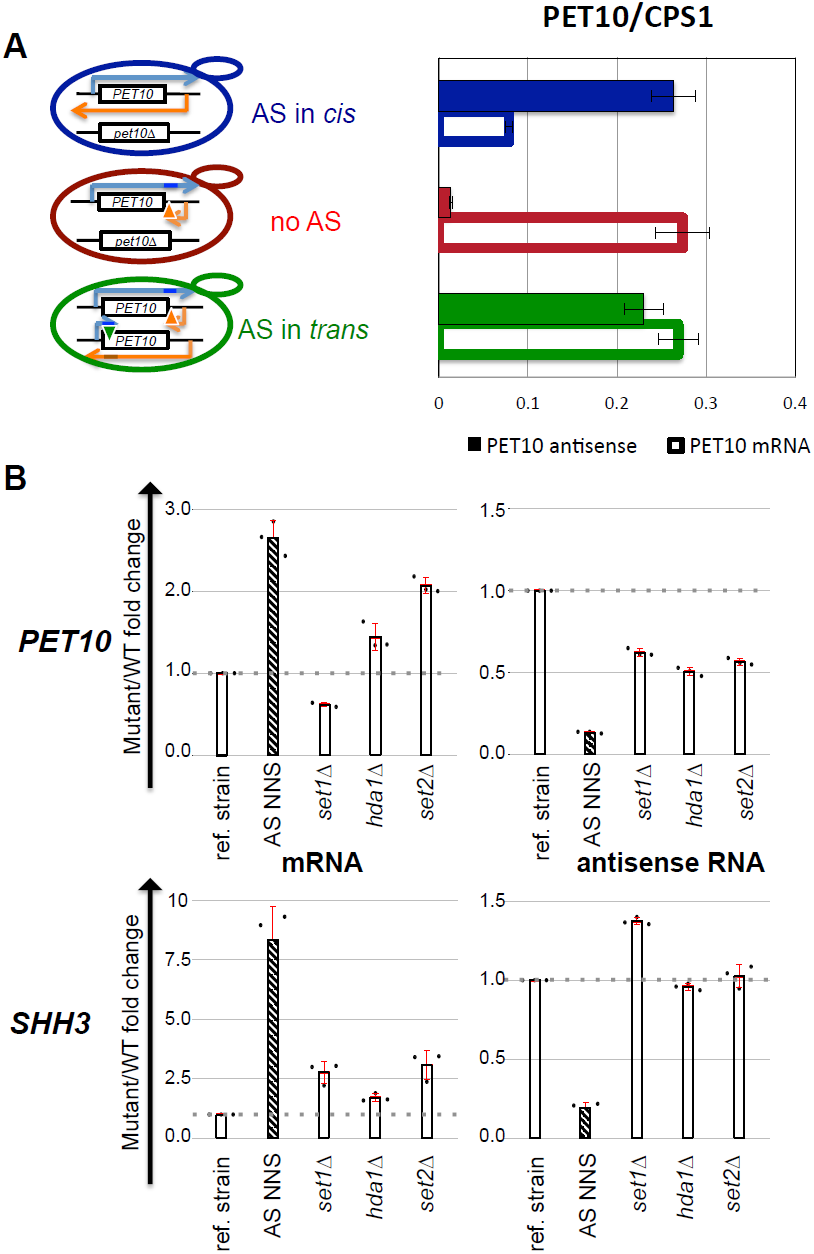
Antisense repression is mediated by transcriptional interference mechanisms. A Strand-specific RT-qPCR analysis of *PET10* mRNA and antisense RNA abundance in diploid strains. PET10 antisense is transcribed *in cis* (blue), in trans (green) or not produced (red). B Strand-specific RT-qPCR analysis analysis of *PET10* (upper panel) and *SHH3* (lower panel) mRNAs and antisense abundances in a mutant strain where the deletion of *UPF1* (ref. strain) is either combined to an antisense NNS terminator insertion (AS NNS, purple; positive control), or to the deletion of a chromatin modification factor (*set1*Δ, *hda1*Δ and *set2*Δ).

### Several PolII elongation-associated chromatin modification factors cooperate to mediate antisense transcriptional interference

As described above, TSS-overlapping asRNA associated genes were more prone to be up-regulated upon deletion of chromatin modifiers such as *SET2* (Figure 2 and Supplementary Figure S3) or *SET1*, *RCO1* and *EAF3* (Supplementary Figure S3, A-D) than the other categories of genes. Although the effects of *set2*Δ, *rco1*Δ and *eaf3*Δ are expected to be largely redundant as these factors act in the same chromatin modification pathway (27, 28, 47), we also observed that more than half (80 out of the 155) of the genes that we computed in the Churchmann dataset (47) as the most up-regulated in *set2*Δ were also up-regulated in *set1*Δ (Supplementary Figure S3E). This suggested that these different chromatin modifiers might cooperate to mediate asRNA transcriptional gene repression. The interpretation of such data are complicated by the fact that chromatin modifiers can affect both the mRNAs and their associated antisense (24, 25). To address this question, we directly measured the effects of *set1*Δ, *set2*Δ and *hda1*Δ on both the asRNA and the mRNA by strand-specific RT-qPCR (see Material and Methods). The analysis of the *PET10* locus shows a complex picture. Not only the mRNAs were positively affected in different mutants, but the levels of asRNA were also impaired in all these mutants, making the evaluation of the antisense transcription interference on the mRNA difficult. In contrast, the *SHH3* asRNA was not repressed in these deletion strains while the mRNA was significantly derepressed in all the mutants, although not at the level of the control strain in which the asRNA transcription elongation is restricted by the NNS-terminator (Figure 6B). This suggests that several chromatin modification pathways cooperate to mediate an efficient transcriptional interference to repress gene expression.

### Sense and antisense transcription can mutually repress each other

As discussed above, the fact that the category of genes associated with asRNAs was enriched in the least expressed genes (bin 1 or quiescence-enriched genes) could result from two non-exclusive phenomena: the asRNA transcription represses the mRNA or the absence of mRNA expression releases pervasive asRNA transcription from 3’-NFRs. We showed that, in four out of five genes tested, asRNA transcription interruption led to an increase of sense mRNA levels, indicating a strong repressive effect of asRNA transcription on mRNA levels. As observed in Figure 4, the expression time courses of the mRNAs and asRNAs present inversed expression patterns, which is compatible with the mutual repression of sense and antisense transcription. We tested this hypothesis by interrupting the mRNA transcription of the *PET10*, *SHH3*, *ARO10* and *MOH1* genes. Figure 7A shows that in all cases but for *MOH1*, restricting transcription elongation led to an up-regulation of the corresponding asRNA at 48 hours (post-diauxic shift). The absence of effect observed for *MOH1* can be explained by the fact that its polyA site is the only one (out of the four genes analysed) located upstream of its associated asRNA TSS (see Supplementary Figure S1). The mutual repression of sense and antisense transcription can thus be observed but is dependent on *locus*-specific architecture.

**Figure 7.**
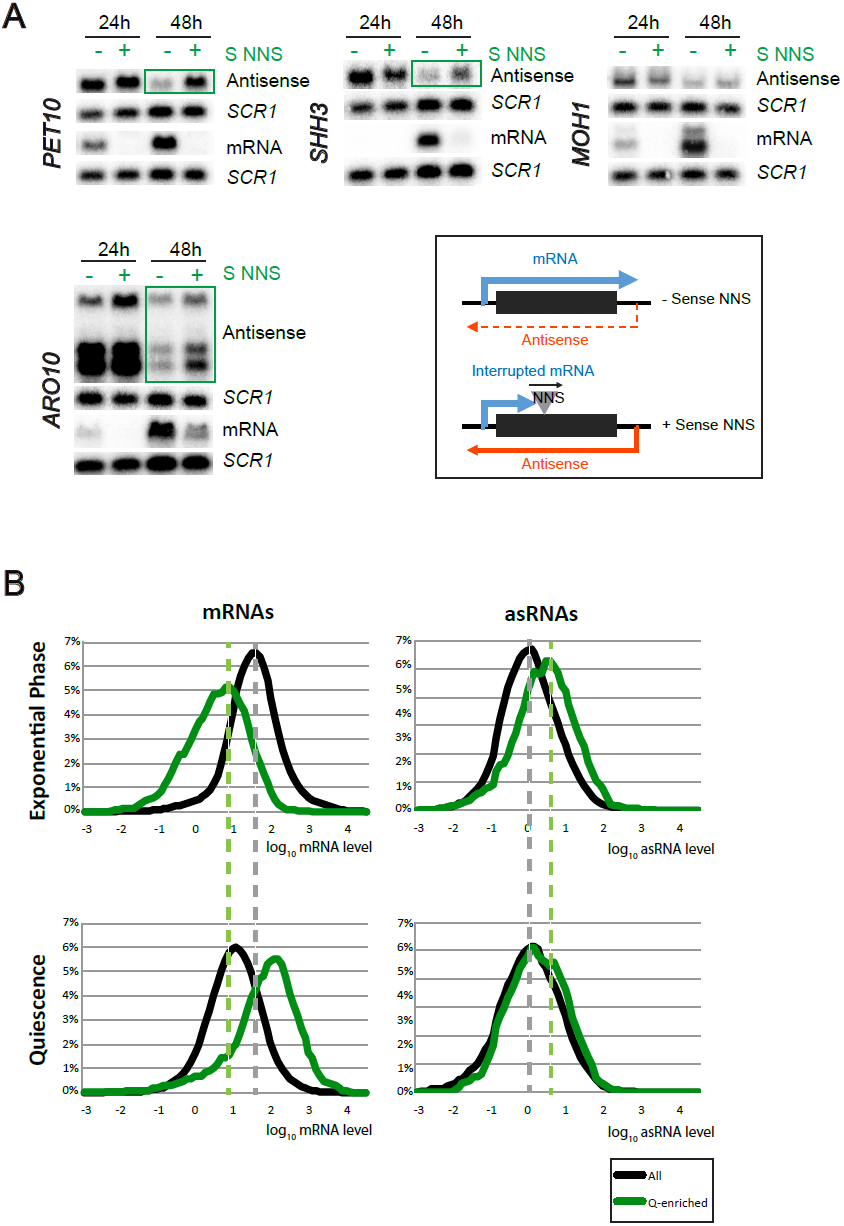
Gene expression is repressive for antisense non-coding transcription. A Northern blot analysis of *PET10*, *MOH1*, *SHH3* and *ARO10* mRNA and antisense RNAs in Δ*upf1* strain, after 24h or 48h of growth in YPD and with (+) or without (−) the insertion of a sense Nrd1-Nab3-Sen1 terminator (NNS S). RNA probes and NNS insertion are described in Supplementary Figure S4 and Supplementary Table S2 (see also Material and Methods for strain construction) and NNS S - corresponding strains in Supplementary Table S1. *SCR1* is used as a loading control. B Comparison of density plots between all (black lines) and Q-enriched genes (green lines) for mRNAs (left panels) or associated asRNA (right panels) from cultures harvested in exponential (upper panels) or G0 (lower panels) phases. Log_10_ RNA levels are plotted (abscissa) function of the frequency (ordinate).

As shown above, the mRNAs and their associated asRNAs exhibited an inverse pattern of expression between the exponential phase and quiescence, as expected if the expression of sense and antisense were mutually exclusive. Q-enriched genes were associated with more antisense transcription than average during the exponential phase, *i.e.* when these genes are repressed. Conversely, one could expect that they would be less associated with asRNA than average in quiescence since they are the genes whose mRNAs are most abundant under this condition. This turned out not to be the case. Indeed, the quiescence-enriched genes remained associated with slightly higher asRNA levels than average even during quiescence (Figure 7B). This is consistent with the observation that there is no obligatory repression of asRNA transcription when sense transcription is induced and with the observation that the quiescence-enriched genes are, overall, more associated with asRNAs than other genes.

The analysis of sense and antisense expression in quiescence also revealed that if the mRNA levels are strongly decreased in quiescence, as expected, the global level of asRNAs did not change markedly (Supplementary Figure S5). A likely explanation is that mRNA transcription interferes with pervasive asRNA transcription during exponential phase. The global repression of transcription in quiescence (50) could then be compensated for the asRNAs reduced interference from mRNA transcription. This was verified at the *HIS1* locus where the strong asRNA observed only in quiescence (Figure 8A) could be revealed during the exponential phase by interrupting the *HIS1* gene transcription by a strand specific NNS terminator insertion (Figure 8B).

**Figure 8.**
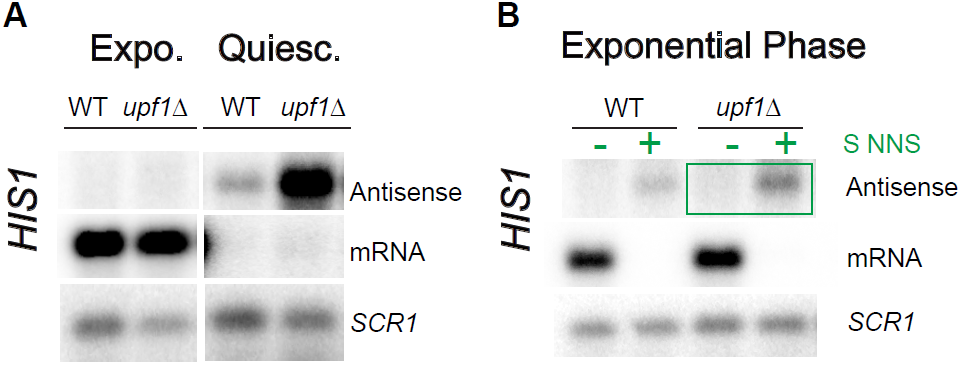
*HIS1* associated asRNA is induced in quiescence or when HIS1 mRNA transcription is interrupted. A, B Northern blot analysis of *HIS1* mRNA and antisense RNAs in WT and Δ*upf1* strains in exponential phase or quiescence (A) or in exponential phase with (+) or without (−) the insertion of a sense Nrd1-Nab3-Sen1 terminator (NNS S; B). RNA probes and NNS insertion are described in Supplementary Figure S1 and Supplementary Table S2 (see also Material and Methods for strain construction) and NNS S-corresponding strains in Supplementary Table S1. *SCR1* is used as a loading control.

## DISCUSSION

Targeted studies have previously described a ten or so of specific examples, in which the transcription of a non-coding RNA was mediating gene regulation (24), see for reviews 51). To what extent antisense-mediated transcription interference affects gene expression genome-wide remains poorly defined. A recent large-scale approach (32), which was used to address this question, only focused on genes associated with asRNAs sufficiently stable to be readily detected in wild type cells (SUTs; 4). It showed that antisense transcription weakly affected the expression of only 12-25% of the SUTs associated genes and no particular class of genes was found to be specifically affected. Here, we addressed the question from a different angle by searching classes of genes frequently presenting characteristics associated with asRNA transcription interference.

In order to identify genes potentially repressed by asRNA transcription interference, we analysed the transcriptome of NMD deficient cells (*upf1*Δ) since abrogating NMD reveals non-coding RNAs normally efficiently degraded by this quality control pathway (13). Using a relatively stringent threshold for antisense detection (see Figure 1C), we defined 816 asRNAs. The less expressed genes were more often associated with asRNA than average and, most interestingly, this bias essentially resulted from a higher number of TSS overlapping asRNAs, reaching 30% of the genes in the bin corresponding to the least expressed genes (bin 1; Figure 1E, Supplementary Figure S2 and Dataset 1). This strongly suggested that antisense mediated transcription interference could contribute to the repression of up to 30% of these least expressed genes (bin 1). Remarkably, genes whose mRNAs were enriched in quiescence relative to exponential growth are mostly found in bin 1 and behaved similarly (Figure 3). It thus defined a family of genes potentially associated with frequent asRNA transcription mediated repression. This hypothesis was strengthened by the observation that genes associated with TSS overlapping asRNAs in bin 1 were also subjected to a regulation by chromatin modification factors much more often than average (Figure 2 and Supplementary Figure S3) and this was particularly true for quiescence-enriched genes (Figures 3E and Supplementary Figure S4B). This was especially noteworthy since asRNA transcription mediated regulation was previously found to affect single genes that belong to diverse genes families.

To test the hypothesis that the full repression of quiescence-enriched genes during the exponential phase often relies on interference by antisense transcription, we directly analysed five of these Q-enriched genes associated with TSS-overlapping asRNA (*PET10*, *CLD1*, *MOH1*, *SHH3* and *ARO10*). For four out of these five genes, specifically interrupting asRNA transcription resulted in a strong induction of the corresponding mRNAs during the exponential phase (Figure 5). Interestingly, the only gene that did not respond was *ARO10* but it was also the least repressed gene in our conditions during the exponential phase (Figure 4). Interestingly, this was the only gene we analysed that was also analysed in the Huber study (32). Consistently with our observations, although they could not find a repressive effect of its associated asRNA in rich medium, they found it to be regulated by antisense transcription when the cells were grown in synthetic complete medium. It thus turns out that the TSS-overlapping asRNAs associated to all five Q-enriched genes we tested can have a repressive role on gene transcription.

If, in a few instance, the asRNA itself was suggested to play a direct role in gene repression, in the majority of cases examined thus far this repressive effect was shown to be mediated in *cis* by antisense transcription interference, the asRNA being only a by-product of this process (see for review 51). Using strand specific NNS terminators in diploid strains, we directly showed, on the *PET10* locus, that the effects we observed act only in *cis*, which confirmed that transcriptional interference is likely the mechanism at play in these examples (Figure 6A).

Chromatin modifiers are though to be key players of transcriptional interference (24). Considering the high redundancy of chromatin modifiers, we can extrapolate that the number of genes submitted to a regulation by them is underestimated (Supplementary Figure S3E). We could effectively measure this redundancy in two examples (*PET10* and *SHH3*). The single deletion of each factor we tested couldn’t reach the complete de-repression that was observed when the asRNA was interrupted (Figure 6B). Taken together, these results strongly suggest that the observed asRNA-transcription mediated repression involves several redundant chromatin modification/remodelling pathways. This is reminiscent of previous observations showing that gene silencing is mediated by redundant mechanisms involving multiple histone modifiers (52).

Interestingly, we found the asRNA repression upon induction of the mRNA to be frequent, although not obligatory and depending on the fact that the induced mRNA transcription overlaps the asRNA TSS (Figure 7 and Supplementary Figure S1). Overall, the asRNA levels remained high in quiescence, even slightly higher than average (Figure 7B, right panels). It suggests a model by which, in contrast to previously studied examples (see for examples 18, 19), the asRNA expression is not regulated by specific transcription regulators. Rather these RNAs would be constitutively expressed, unless repressed by sense transcription when mRNAs are induced and overlap their TSSs. Their transcription would thus act “passively” as an amplifier of gene regulation, turning an non-induction into repression, as previously suggested for the SUR7 gene (29). Consistent with this model, blocking asRNA transcription elongation during the exponential phase resulted in a 2.6 and 8.3 fold increase of *PET10* and *SHH3* mRNAs respectively (Figure 6B), which is markedly lower than the induction estimated by comparing the increase of their relative expression levels measured from the quiescence versus exponential phase transcriptome datasets (5.9 and 264.6 fold increase respectively in the *upf1*Δ background; Dataset 1).

In our study, we demonstrated that TSS-overlaping antisense-mediated transcriptional interference is a frequent mechanism used for full gene repression. This mechanism is often hidden since these antisense transcripts are rapidly degraded by the NMD pathway and therefore not detected in wild type conditions.

Making more complex the overall picture, we and others reported the existence of conditional asRNAs, such as for example the *HIS1* antisense RNA specifically expressed during G0 (Figure 8A), or Meiotic Unannotated transcripts (MUTs; 53). In addition, asRNAs were shown to mediate protein expression regulation depending on various growth conditions (32). Widespread antisense transcription has thus the potential to repress the synthesis of sense RNA and participate to differential gene expression and adaptation to various environmental and growth conditions.

Interestingly, the presence of a PET10 asRNA with an inversed expression profile compared to the mRNA was shown to be conserved in all five analysed *Saccharomyces* species, supporting its functional role (54). More generally, a phylogenetic conservation study of lncRNAs in budding yeasts has shown that, since the divergence with *N. castellii*, which has retained a functional RNAi machinery, the level of asRNAs and their extent has globally increased. Accordingly, this suggested that the lack of RNAi favoured the development of asRNA transcription mediated gene regulation (55).

Revealing the importance of antisense pervasive transcription and its interplay with gene expression, our study highlighted for the first time the importance of antisense-mediated transcriptional interference and the mutual repression on gene versus antisense transcription depending on growth conditions. We could estimate that this mechanism concerned up to 30% of the least expressed genes and resulted in a strong and efficient gene repression.

## ACKNOWLEDGEMENT

We thank Frank Feuerbach for providing strains LMA2811 and LMA2819. We thank Bernard Turcotte, Cosmin Saveanu and Micheline Fromont-Racine for discussions and critical reading of the manuscript. We acknowledge Jean Yves Coppee and Caroline Proux for the facilities and expertise of the Transcriptomic Platform (PF2) for RNAseq experiments.

## DATA AVAILABILITY

The data reported here have been deposited in NCBI GEO under the accession number GSE101368.

